# Kaposi’s sarcoma-associated herpesvirus glycoprotein K8.1 is critical for infection in a cell-specific manner and functions at the attachment step on keratinocytes

**DOI:** 10.1101/2023.03.19.533316

**Authors:** Shanchuan Liu, Anna K. Großkopf, Xiaoliang Yang, Stefano Scribano, Sarah Schlagowski, Armin Ensser, Alexander S. Hahn

**Author notes:** Address correspondence to Alexander S. Hahn,. HIV & AIDS Malignancy Branch, Center for Cancer Research, National Cancer Institute, Bethesda, Maryland, United States.

## Abstract

Kaposi’s sarcoma-associated herpesvirus (KSHV) is associated with Kaposi’s sarcoma and several B cell malignancies. K8.1, the major antigenic component of the KSHV virion, has been reported to play a critical role in the infection of certain B cells, but otherwise its function remains enigmatic. We created a K8.1 knockout virus (KSHVΔK8.1) in the BAC16 genetic background and analyzed its infectivity on a range of adherent cells. We observed a strong defect on several epithelial cells, e.g. the HaCaT keratinocyte model cell line, HEK 293T and A549 lung epithelial cells, but no such defect on other cells, among them e.g. lymphatic and blood endothelial cells. Mechanistically, we found that reduced infectivity of the K8.1 knockout virus correlated with reduced attachment to HaCaT cells. The defect in infectivity of KSHVΔK8.1 could be rescued by complementation through expression of K8.1 in KSHVΔK8.1 producing cells by means of a lentiviral vector. In a coculture infection model, KSHVΔK8.1 was highly efficient at infecting the BJAB B cell line but was significantly impaired at infecting the MC116 B cell line, in line with a previous report. In fusion assays together with the gH/gL glycoprotein complex and gB, the components of the conserved herpesviral core fusion machinery, we did not observe activation of membrane fusion by K8.1 or its R8.1 homolog of the rhesus monkey rhadinovirus. In summary, we found K8.1 to function in a highly cell-specific manner during KSHV entry at the attachment step, playing an important role in the infection of epithelial cells.

**IMPORTANCE:** Kaposi’s sarcoma-associated herpesvirus (KSHV) is the causative agent of several B cell malignancies and Kaposi’s sarcoma. We analyzed the function of K8.1, the major antigenic component of the KSHV virion, in the infection of different cells. To do this, we deleted K8.1 from the viral genome. It was found that K8.1 is critical for the infection of certain epithelial cells, e.g. a skin model cell line, but not for infection of many other cells. K8.1 was found to mediate attachment of the virus to cells where it plays a role in infection. In contrast, we did not find K8.1 or a related protein from a closely related monkey virus to activate fusion of the viral and cellular membranes, at least not under the conditions tested. These findings suggest that K8.1 functions in a highly cell-specific manner during KSHV entry, playing a crucial role in the infection of e.g skin epithelial cells at the attachment step.

## INTRODUCTION

KSHV is the etiological agent of Kaposi’s sarcoma (KS) and is associated with at least two B cell malignancies, primary effusion lymphoma and a variant of multicentric Castleman’s disease. An association with KSHV-positive diffuse large B cell lymphoma was also reported (1–5). A recent report suggests association with osteosarcoma (6), and the virus is associated with the so-called KSHV inflammatory cytokine syndrome (7). KSHV-associated malignancies constitute a major disease burden in Sub-Saharan Africa and in immunocompromised individuals worldwide, in particular in the context of HIV infection (reviewed in (8, 9)). In some African countries KS is the cancer that is associated with both the highest morbidity and mortality among all types of cancer (10). A vaccine to prevent either KSHV primary infection or the development of KSHV-associated diseases would be highly desirable. For both approaches, understanding the function of the individual glycoproteins at the different stages of KSHV entry into specific host cells is critical to understand their role during host colonization of individual tissues.

The roles of the individual glycoproteins (GPs) of KSHV during the different steps of the entry process, attachment, endocytosis, and membrane fusion, have so far only been poorly defined in the context of different types of host cells. Like other herpesviruses KSHV encodes the glycoproteins gB, gH, and gL that together form the conserved core fusion machinery of the herpesviruses (11). While gB is the fusion executor, the interaction of KSHV gB with integrins through an RGD motif may rather contribute to attachment or endocytosis (12–15). The interaction of gH/gL with receptors from the Eph family can trigger membrane fusion by the core fusion machinery (16, 17), similar to what is observed with the related human Epstein-Barr virus or the rhesus monkey rhadinovirus (16, 18). This at least holds true within the experimental limits of these reports as gB always had to be substituted for a gB of a related virus, either EBV or RRV, due to KSHV gB’s extremely low fusogenicity, which makes it unsuitable for assays measuring cell-cell fusion. Engagement of Eph family receptors also triggers endocytosis (19–21). The exact function and molecular interactions of a proposed fusion receptor, xCT, which normally forms a complex with CD98, so far remain unclear, with different reports claiming either a direct role in fusion or for post-entry gene expression (15, 22).

The role of KSHV glycoprotein K8.1, also termed K8.1A or gp35-37 (23), is not well understood. It binds heparan sulfate (24) and is the major antigenic component of the KSHV virion (23). A role of K8.1, that is independent from its binding to heparan sulfate, in the infection of the MC116 B cell line as well as in the infection of primary tonsillar B cells has been described (25). Otherwise, its function remains enigmatic. For example, it is not known whether K8.1 plays a role in the fusion process, something that could be suspected due to its potential positional homology to the gp42-encoding gene of EBV within the KSHV genome. EBV gp42 clearly functions as a trigger of fusion (26). Another question is whether K8.1 only plays a role for infection of some B cell populations or also for infection of other cell types. We therefore generated a K8.1 deletion mutant and analyzed its relative infectivity on different primary cells and cell lines. We also investigated K8.1’s contribution to virus attachment and triggering of the fusion machinery.

## RESULTS

### Deletion of K8.1 results in decreased infectivity on several cell lines of epithelial origin

Previously, a role for K8.1 in the infection of B cells was reported by Dollery et al. (25) but a comprehensive analysis of infection in other cell types was not performed in that study. We therefore constructed a K8.1 deletion mutant, with a mutation/deletion of the start codon and signal peptide of K8.1. While probably not necessary for this *in vitro* study, the strategy of introducing a larger deletion in the signal peptide region was chosen to make sure that no alternative start codon can give rise to a functional protein and to prevent reversion, also e.g. in an experimental animal host in future studies. The deletion in the K8.1 locus (Fig. 1A) resulted in the complete absence of the typical staining pattern with anti-K8.1 antibody observed after Western blot analysis of virus preparations (Fig. 1 B, leftmost lanes) or cell lysates (Fig. 1 B, rightmost lanes) from iSLK cells that had been previously transfected to harbor BAC 16 wt or ΔK8.1 and were induced to enter the lytic cycle. Whether the single band slightly above 100kDa, which is recognized by the anti-K8.1 monoclonal antibody, represents a background band or a K8/K8.1 fusion transcript (27) is at present not entirely clear. We next compared wt and ΔK8.1 virus on a range of different cell lines and primary cells (Fig. 2 A). While infection of the SLK cell line (a clear renal carcinoma cell line) (28), U2197 (histiocytoma cell line), and of primary human foreskin fibroblasts (HFF), human umbilical vein endothelial cells (HUVEC), and lymphatic endothelial cells (LEC) cells was comparable, the ΔK8.1 virus exhibited a significant defect on 293T cells and on A549 cells. On HaCaT cells, the defect was also very pronounced but due to the overall low susceptibility of HaCaT could only be shown at higher MOI after spin infection. The discrepancy in susceptibility of HaCaT to infection by wt and ΔK8.1 virus in comparison to SLK cells was also confirmed in separate experiments with another pair of virus stocks (Fig. 2 B), and it was not dependent on e.g. the calcium concentration in the culture medium, which reportedly controls differentiation of HaCaT (29). KS is primarily a cutaneous tumor and skin keratinocytes are infected by KSHV, also in the context of KS lesions (30–32). HaCaT are a model cell line for keratinized epithelia that retain the features of primary skin keratinocytes (33), and we therefore focused our studies on this cell line.

**Figure 1.**
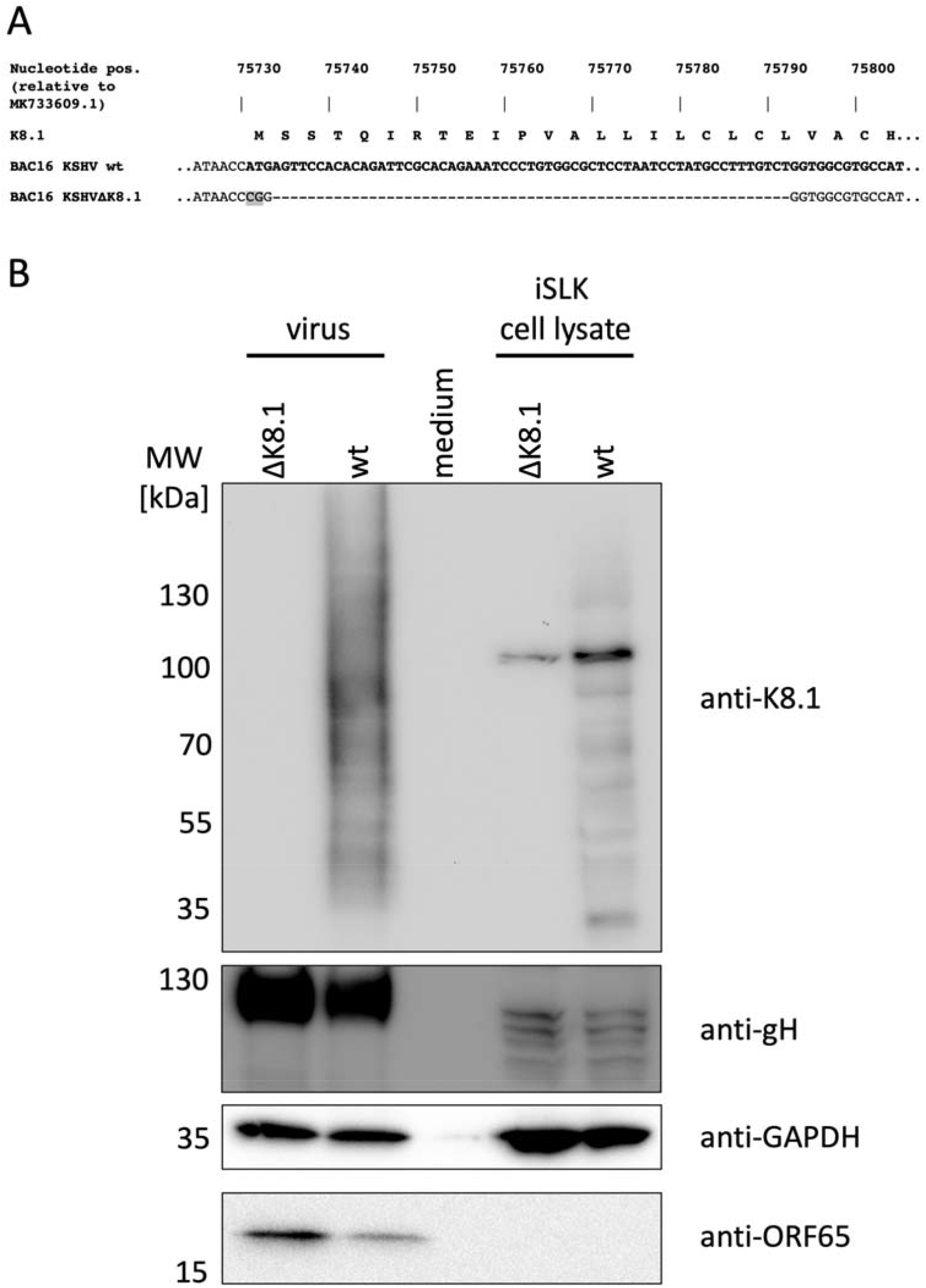
Construction of KSHVΔK8.1. A) Mutation and deletion of the K8.1 start codon and signal peptide in the BAC16 genome. B) Loss of the characteristic high-molecular-weight forms of K8.1 in virus preparations of KSHVΔK8.1 or iSLK producer cell lysate as detected by Western blot. Cell culture medium serves as a background control.

**Figure 2.**
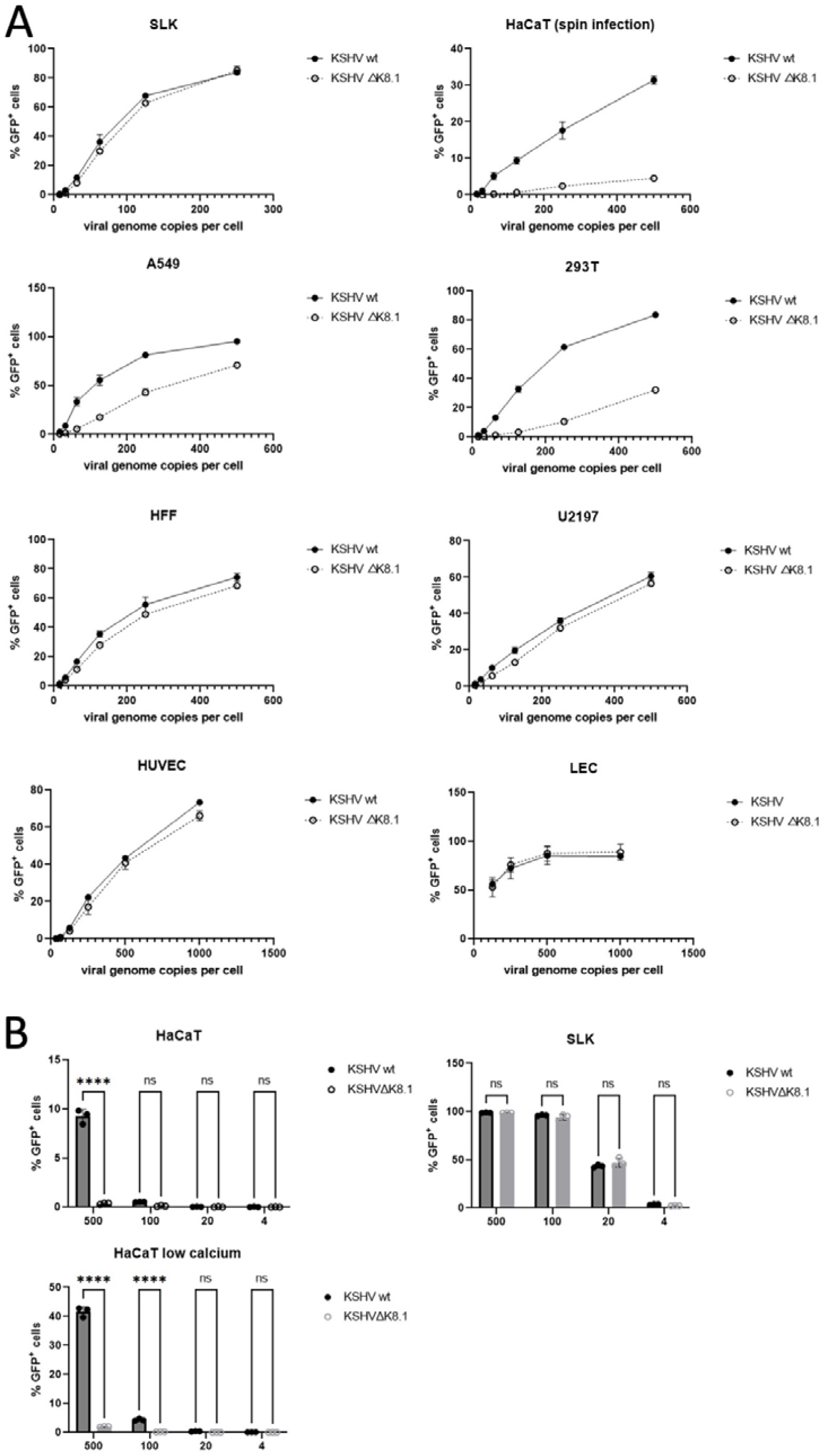
Loss of K8.1 results in cell-specific defects in infectivity on different types of adherent cells. A) Different cell lines and primary cells were infected with BAC16-derived KSHV wt or KSHVΔK8.1 using the same number of virus particles as measured by DNAse-resistant genome copies. Infection was quantified by the percentage of GFP reporter gene-expressing cells. One representative experiment of two independent experiments with two independent pairs of virus stocks is shown. B) Repeat experiment comparing infection of SLK and HaCaT (at normal cell culture conditions and at low calcium) by KSHV wt and KSHVΔK8.1. **** = p<0.0001, ns = non-significant. Two-Way ANOVA with Sidak’s correction for multiple comparisons, n=3.

In order to control for possible non-specific off-target effects on viral infectivity that may result from our genetic manipulation of the K8.1 locus, we performed a reconstitution experiment. iSLK cells harboring BAC16ΔK8.1 were transduced with either empty lentiviral vector (control) or a lentiviral vector coding for Tandem-Strep-tagged K8.1. We then compared infection by KSHV wt or KSHVΔ8.1 produced from cells either transduced with empty lentiviral vector or with lentiviral vector encoding K8.1. K8.1-reconstituted KSHVΔK8.1 contained K8.1 in the virus preparation (Fig. 3 A) and infectivity of KSHVΔK8.1 on HaCaT, A549 and 293T cells was visibly rescued by reconstitution of K8.1 expression (Fig. 3 B). It should be noted that lentiviral transduction may not lead to uniform expression of the transgene in all cells to the same degree and may be limited with regard to the achievable expression levels, which likely precludes complete reconstitution, as K8.1 is expressed to high levels during lytic KSHV replication.

**Figure 3.**
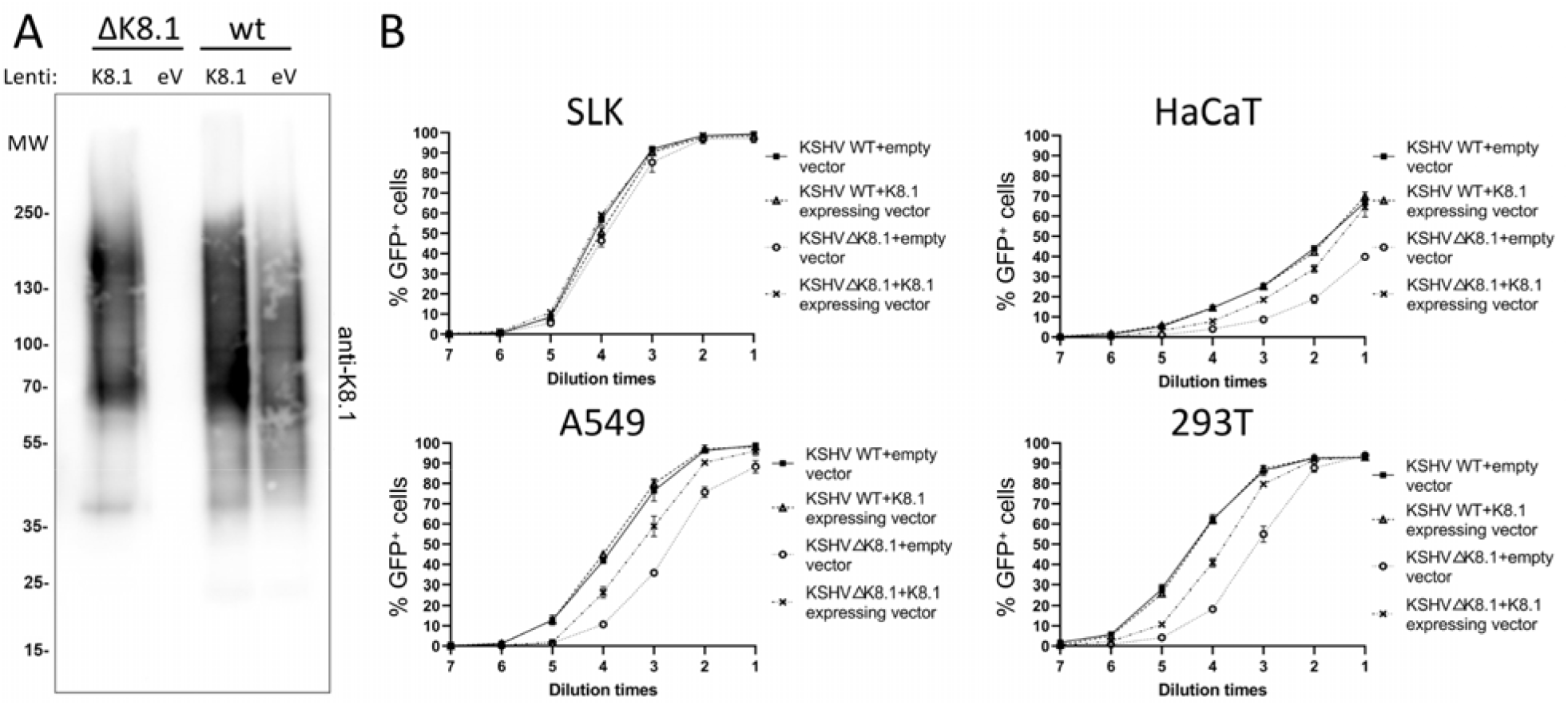
Recombinant K8.1 expression in virus producing cells reverts the cell-specific defect in infectivity of KSHVΔK8.1. A) Western blot analysis of virus preparations from chemically induced iSLK producer cells harboring BAC16 KSHVΔK8.1 or wt, transduced with empty vector or a lentiviral vector coding for K8.1 tagged with a C-terminal tandem StrepTag. (Lenti = lentiviral vector; eV = empty vector) B) Infection of SLK, HaCaT, A5459, and 293T. HaCaT were infected by spin infection. Error bars represent the standard deviation of triplicate values. The data show one representative experiment out of two independent experiments.

### Deletion of K8.1 results in reduced cell-to-cell transmission into the MC116 B cell line but not into the BJAB or Raji B cell lines

Our previous work demonstrated that the BJAB B cell line is infected in a manner that is highly dependent on the EphA7 receptor and to a lesser degree on the EphA5 receptor, and that both receptors are engaged by the KSHV gH/gL complex, similar to the interaction with the high-affinity EphA2 receptor (34). Work by Dollery and colleagues has demonstrated a critical role of K8.1 for the infection of the MC116 B cell line (25). As neither of the two cell lines in our hands supported infection with free virus (not shown), which disagrees with previous results for MC116 (35) but may be rooted in e.g. different culture conditions that lead to differences in susceptibility, we could not compare the two viruses as free virus on those cells. We instead compared both cells lines head to head in our established cell-to-cell-transmission model for KSHV into B cell lines (34) that roughly follows the established protocol by Myoung et al (36), who first described the critical role of cell-to-cell transmission for infection of lymphoblastoid B cell lines. We found that in agreement with Dollery et al. (25) MC116 cells infection was highly dependent on K8.1, even if MC116 in our hands exhibited comparatively low susceptibility (Fig. 4 middle column pair). On the other hand, infection of BJAB or Raji was completely independent of K8.1 (Fig. 4 left and right column pairs), again demonstrating that the deletion in KSHVΔK8.1 does not cause a general defect in infectivity or induction of the lytic cycle.

**Figure 4.**
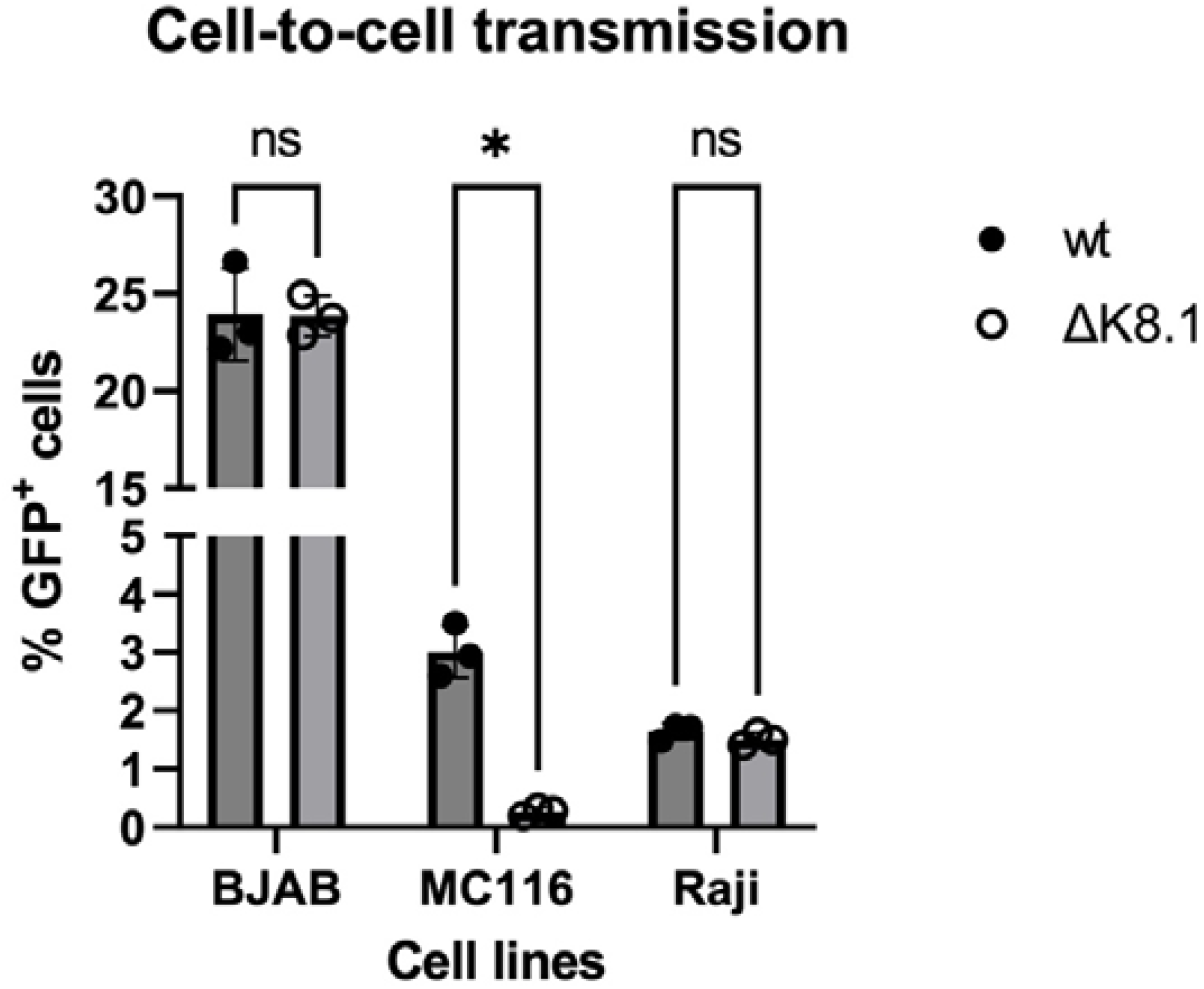
KSHVΔK8.1 exhibits a defect in cell-to-cell transmission into the MC116 B cell line. iSLK cell carrying the BACs coding for the respective viruses were chemically induced to enter the lytic cycle and were co-cultured with the indicated target cells. Infection was determined after 4 days by flow cytometry. Error bars represent the standard deviation;*: p<0.05, 2-way-ANOVA with Sidak’s correction for multiple comparisons, n=3.

### Deletion of K8.1 results in reduced attachment to HaCaT skin epithelial cells

The defect in infectivity that was observed with KSHVΔK8.1 on HaCaT cells could conceivably be caused at different stages. As K8.1 is a glycoprotein, the attachment step, endocytosis of the virus particle or fusion of viral and cellular membranes come to mind. We analyzed attachment by replicating the spin infection protocol used for the infection assay but immediately after centrifugation at 4° C, the cells were washed with cold PBS and then harvested for quantification of cell-associated, DNAse-resistant viral nucleic acid as a surrogate marker for attachment. We found that attachment of KSHVΔK8.1 to HaCaT cells was clearly and significantly reduced by over twofold in comparison to attachment of KSHV wt (Fig. 5 A).

**Figure 5.**
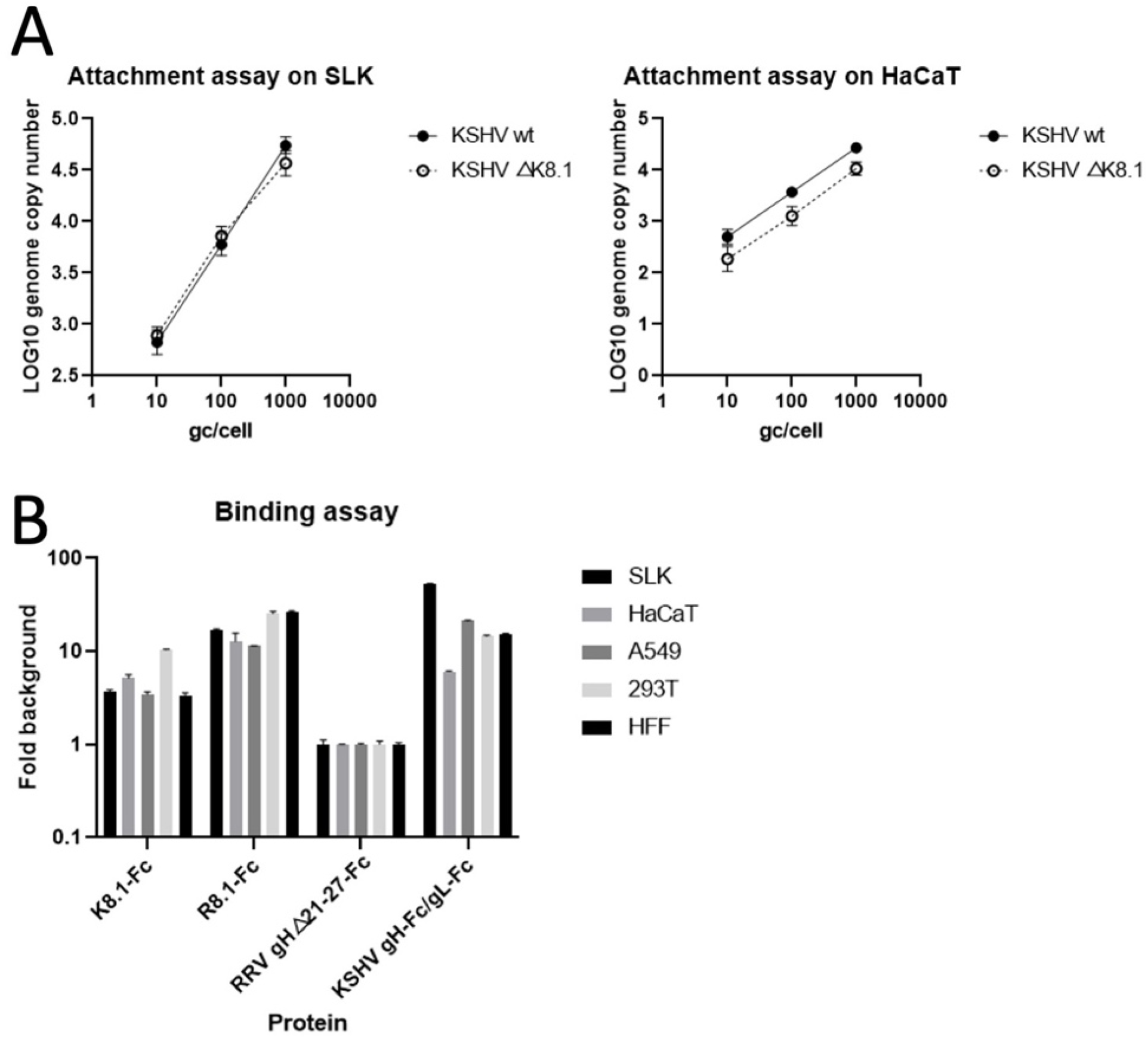
KSHVΔK8.1 exhibits reduced attachment on HaCaT cells. A) SLK cell or HaCaT cells were spin-infected at 4° C with the indicated amounts of virus and bound viral DNA was quantified by qPCR. Binding curves for KSHV wt and KSHVΔK8.1 were indistinguishable on SLK but significantly different for HaCaT. (p=0.0051, Non-linear fit for semi-log data by least squares regression using mean values for each data point, n=6) B) Binding of Fc-fusion proteins K8.1-Fc, gH-Fc/gL, R8.1-Fc or RRV gHΔ21-27-Fc (control) to the indicated cells was measured by flow cytometry. Error bars represent the standard deviation, n=3.

### The importance of K8.1 for infection is partially reflected in its binding to cellular membranes

In order to determine whether K8.1’s function is related to its quantitative binding to the membrane of target cells, which might mediate attachment, or rather an indirect effect of its presence in the virion, we performed binding assays with a K8.1-Fc fusion protein that consists of the extracellular domain of K8.1 fused to the Fc part of immunoglobulin G (Fig. 5 B). For comparison, we also used similar molecules, gH-Fc/gL of KSHV that binds to heparan sulfate and Eph family receptors (19, 37), and as control RRV gH-Fc mutated in the Plxdc1/2 receptor binding site and unable to bind Eph family receptors without gL (RRV gHΔ21-27-Fc) (18), a molecule that in our hands does not bind to any cell tested so far. Binding of K8.1-Fc measured as fold mean fluorescence intensity over control was clearly most pronounced on 293T cells, followed by HaCaT, two cell lines where the most pronounced defect of KSHVΔK8.1 was observed. For HaCaT, in addition to high K8.1 binding, very low binding of gH-Fc/gL was observed, in line with low expression of EphA2 in this cell line as reported by the human protein atlas (https://www.proteinatlas.org/). For A549, the third cell line where KSHVΔK8.1 showed a clear-cut defect, the picture was less clear. A549 exhibited K8.1 binding that was comparable to that on other cells and also interacted avidly with gH-Fc/gL. Therefore, the relative importance of K8.1 for infection may be reflected by the abundance of cellular interaction partners like heparan sulfate proteoglycans on the cell surface, but that does not entirely explain its role.

### Presence or absence of K8.1 or its RRV homolog does not appreciably modulate gH/gL-activated membrane fusion by gB

While our results so far pointed to a function of K8.1 at the attachment step, mediating binding of the virus to host cell membranes, a function in fusion is also possible. The EBV glycoprotein gp42 for example, which is located at a similar or even the same position in the EBV genome compared to K8.1 in the KSHV genome, triggers fusion (26). As KSHV gB is barely fusogenic in cell-cell fusion assays, analyzing the trigger function of gH/gL on gB-mediated membrane fusion is usually carried out using a gB from a related virus (17, 38). As established in our group, we tested KSHV gH/gL together with gB of the related rhesus monkey rhadinovirus. Addition of K8.1 to this combination of glycoproteins did not result in an increase in fusion of effector cells expressing gH/gL/gB or gH/gB (Fig. 6 A). We tested fusion activity with and without gL, which results in the ability or inability to use Eph family receptors for fusion (39). This approach might demonstrate modulation of fusion that might be masked by Eph-mediated membrane fusion in the presence of the full gH/gL complex. Even so, under none of these conditions did K8.1 modulate fusion activity. As this only demonstrates a lack of action on gH/gL-mediated fusion and due to the heterologous gB in the assay, direct activation of gB by K8.1 might be impaired without any obvious effect in our assay if one postulates that R8.1 may not interact with KSHV gB as K8.1 does. We therefore tested the positional homolog R8.1 of the related RRV together with its cognate gH/gL, and gB. Similar to our results with KSHV K8.1, we did not observe appreciable activation of cell-cell fusion on a range of different target cells by RRV R8.1 (Fig. 6 B).

**Figure 6.**
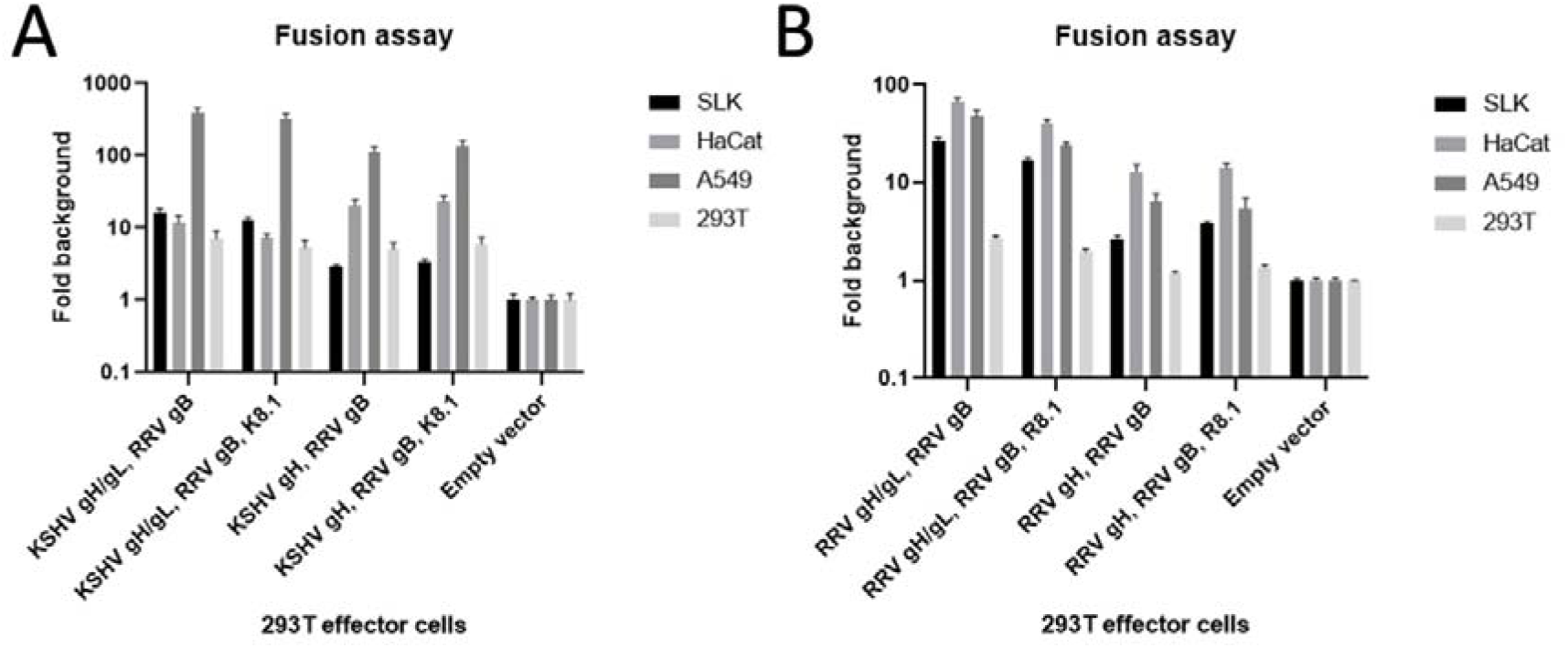
Presence or absence of K8.1 or its RRV homolog do not appreciably modulate gH/gL-activated membrane fusion by gB. A) KSHV gH/gL together with RRV gB or KSHV gH together with RRV gB were expressed in the presence or absence of K8.1 on 293T effector cells, and these cells were co-cultured with the indicated target cell populations. Errors bars represent the standard error of the mean, n=15. B) RRV gH/gL together with RRV gB or RRV gH together with RRV gB were expressed in the presence or absence of R8.1 on 293T effector cells, and these cells were co-cultured with the indicated target cell populations. Errors bars represent the standard error of the mean, triplicate values.

## DISCUSSION

Taken together, our results demonstrate that in addition to the known function of K8.1 in the infection of B cells it has a critical function in the infection of epithelial cells, as observed here for A549, 293T, and the skin model HaCaT cell line, which is not-derived from a tumor but from a spontaneous immortalization event of cultured skin cells and is regarded as highly representative of skin epithelium (33, 40). The specificity of the ΔK8.1 phenotype for certain cells is striking as the K8.1 deletion had absolutely no effect on the infectivity of KSHV on primary fibroblasts, primary endothelial cells and on the SLK and U2197 cell lines. The lack of a phenotype of KSHVΔK8.1 on primary endothelial cells is in agreement with a previous report on the function of K8.1 (25).

While we found a clear correlation of the K8.1 deletion with reduced attachment as demonstrated for HaCaT cells, we also found reduced infection of MC116 cells in a cell-to-cell transmission setting, but normal infection of BJAB cells and Raji cells (Fig. 4). This would be compatible with the findings by Dollery et al., who described inhibition of infection of primary tonsillar B cells by K8.1-targeting antibodies but could not achieve a complete block, unlike as with MC116 cells (25), suggestive of both K8.1-dependent and K8.1-independent infection also in primary B cell preparations. These previous findings and the difference between the BJAB, Raji, and MC116 cell lines argue for the existence of more than one single mechanism for the infection of B cells, which constitute a diverse population.

Our findings in the cell-to-cell transmission setting also suggest a function of K8.1 that goes beyond the canonical concept of attachment as cell-to-cell transmission is believed to function through some form of transfer of infectious virus from one cell to the other or even through a so-called virological synapse, which conceivably might obviate the need for the virus to attach to the target cell, even if that is so far speculation for KSHV. Nevertheless, this notion would fit with the report by Dollery et al. that K8.1’s function for infection of MC116 is independent of heparan sulfate binding (25), which is generally believed to play a role for attachment of viruses to cell membranes. On the other hand, we so far have no indication that K8.1 or its close homologue, R8.1 of RRV, can trigger the core fusion machinery. Therefore, the exact mechanism how K8.1 drives infection remains to be further elucidated. There is also the possibility that while attachment in the original sense is not required during cell-to-cell transmission, the “hand-over” of virus from one cell to the other still requires strong binding to the target cell. In light of our results and previous reports (25), it is conceivable that K8.1 acts through both its interaction with heparan sulfate proteoglycans and through additional interactions that may play a comparatively more important role during infection of B cells or A549 cells, that did not show increased K8.1 binding (Fig. 5 B). It should also be noted that heparan sulfate proteoglycans contain a highly complex mixture of sulfated sugars that may differ between cells (reviewed in (41)) and that the preferred heparan sulfate moieties can be quite specific, as was also observed for gH of KSHV (37).

K8.1 may not only mediate attachment to epithelial cells but also aid in endocytosis of the virus e.g. into MC116 cells, which would be compatible with our findings in adherent cells and in the cell-to-cell transmission model for B cell lines. In this scenario, K8.1 would function more analogous to EBV gp350, which mediates binding and endocytosis, particularly during B cell infection (42), than to EBV gp42, which triggers membrane fusion (26).

In summary, we not only confirm K8.1’s role for the infection of B cells as exemplified by the MC116 line, but we also demonstrate a critical function during infection of several epithelial cells, among them skin-derived keratinocytes. Ultimately, animal experiments will be needed to faithfully elucidate the role of individual glycoproteins for cellular tropism and establishment of persistent infection. With regard to the development of a glycoprotein-based KSHV vaccine, our results argue for the inclusion of K8.1 in such an approach as K8.1 clearly functions during infection of B cells and of skin-derived keratinocytes. Skin keratinocytes have previously been reported to be susceptible to KSHV and to undergo a certain degree of immortalization through infection (31), and skin is an organ that is prominently affected by KS (8). Epidermal keratinocytes were shown to express cyclin D mRNA in cutaneous KS lesions (30) and were shown to be KSHV-infected in such lesions (32), indicating that keratinocytes are at least to some degree involved in cutaneous KS. The role of cutaneous infection in KSHV transmission is unclear. Overall, in light of our findings here and previous results (19, 34, 43), targeting the gH/gL complex to block binding to Eph family receptors together with K8.1 to selectively inhibit infection of certain B cell populations (25) and impair infection of epithelial cells might be a good starting point for the design of vaccine antigens.

## MATERIAL AND METHODS

### Cells

SLK cells (NIH AIDS Research and Reference Reagent Program), A549 (laboratory of Stefan Pöhlmann, German Primate Center-Leibniz Institute for Primate Research, Göttingen, Germany), Human embryonic kidney (HEK) 293T cells (laboratory of Stefan Pöhlmann), HaCaT human keratinocytes (RRID: CVCL_0038), human foreskin fibroblasts (HFF) (laboratory of Klaus Korn, Universitätsklinikum Erlangen, Institute for Clinical and Molecular Virology, Erlangen, Germany), U2197 (Leibniz Institute DSMZ-German Collection of Microorganisms and Cell Cultures GmbH) were cultured in Dulbecco’s Modified Eagle Medium (DMEM), high glucose, GlutaMAX, 25mM HEPES (Thermo Fisher Scientific) supplemented with 10% fetal calf serum (FCS) (Thermo Fisher Scientific), and 50μg/ml gentamicin (PAN Biotech). For low calcium conditions, DMEM high glucose without calcium was supplemented as above and brought to 0.03 mmol calcium from a 2.5 M stock solution. iSLK Bac16 cells (44, 45) (a kind gift of Jae Jung and Don Ganem) were maintained in DMEM supplemented with 10% FCS, 50μg/ml gentamicin, 2.5μg/ml puromycin (InvivoGen), 250μg/ml G418 (Carl Roth) and 200ug/ml hygromycin (InvivoGen). Human vascular endothelial cells (HUVEC) (PromoCell) were maintained in standard Endothelial Cell Growth Medium 2 (PromoCell). Human lymphatic endothelial cells (LEC) (the purchased one) were maintained in Endothelial Cell Growth Medium MV 2 (PromoCell). Bjab (Leibniz-Institut DSMZ-Deutsche Sammlung von Mikroorganismen und Zellkulturen GmbH) and MC116 (a kind gift from Frank Neipel) were cultured in RPMI 1640 medium (Thermo Fisher Scientific) supplemented with 10% fetal calf serum (FCS) and 50 g/ml of gentamicin (R10 medium).

### Deletion of K8.1 in BAC16

The KSHV K8.1 deletion mutant (KSHVΔK8.1) was generated using a two-step, markerless λ-red-mediated BAC recombination strategy as described by Tischer et al. (46). KSHVΔK8.1 harbors a mutation of the K8.1 start codon to CGG followed by a 59-nucleotide deletion resulting in an additional frameshift. In short, the recombination cassette was generated from the pEPKanS template by polymerase chain reaction (PCR) with Phusion High Fidelity DNA polymerase (Thermo Fisher Scientific) using long oligonucleotides (Ultramers; purchased from Integrated DNA Technologies (IDT)) (KSHV_K8.1Mut_F: TAAAGGGACCGAAGTTAATCCCTTA*ATCCTCTGGGATTAATAACCCGGGGTGGCGTGCCATGCCAAT TGTCCCACGTATCG*<u>AGGATGACGACGATAAGTAGGG</u>, KSHV_K8.1Mut_R: CTCTTGCCAGAATCCCAAATGCGAA*CGATACGTGGGACAATTGGCATGGCACGCCACCCCGGGTTAT TAATCCAGAGGAT*<u>CAACCAATTAACCAATTCTGATTAG</u>). The recombination cassette was transformed into BAC16-carrying *Escherichia coli* strain GS1783, followed by kanamycin selection, and subsequent second recombination under 1% L(+)arabinose (Sigma-Aldrich)- induced I-SceI expression. Colonies were verified by PCR of the mutated region followed by sequence analysis (Macrogen), pulsed-field gel electrophoresis and restriction fragment length polymorphism. For this purpose, bacmid DNA was isolated by standard alkaline lysis from 5ml liquid cultures. Subsequently, the integrity of bacmid DNA was analyzed by digestion with restriction enzyme *Xho*I and separation in 0.8% PFGE agarose (Bio-Rad) gels and 0.5×TBE buffer by pulsed-field gel electrophoresis at 6 V/cm, 120-degree field angle, switch time linearly ramped from 1s to 5s over 16 h (CHEF DR III, Bio-Rad). Infectious KSHV recombinants were generated by transfection of purified bacmid DNA (NucleoBond Xtra Midi (Macherey-Nagel)) into iSLK cells using GenJet Ver. II (Signagen) according to manufacturer’s instructions. After visible GFP expression, selection was performed using 200μg/ml hygromycin B (InvivoGen) until only GFP positive cells remained. Lytic replication of KSHV-BAC16 was induced in DMEM supplemented with 10% fetal calf serum (FCS) and 50μg/ml gentamicin by addition of 2.5mM sodium-butyrate and 1μg/ml doxycycline. Supernatant was harvested after the cell monolayer was destroyed. The BAC sequence was verified by Illumina-based next generation sequencing.

### Recombinant virus production and Western blot

iSLK Bac16 WT or ΔK8.1 cells were transduced with lentivirus harboring K8.1 overexpressing vector. After selection by blasticidin (InvivoGen), cells were induced by 2.5 mM NaB and 1 μg/ml doxycycline for 3 days. After that, the produced virus was harvested and subjected to Western blot analysis after concentration by centrifugation through a 5% Optiprep cushion as described previously (47). Western blot analysis was performed as described previously (47), ORF65 was detected using a cross-reactive mouse monoclonal antibody to RRV ORF65 (a kind gift from Scott Wong). K8.1 was detected using monoclonal antibody BS555 (48), a kind gift from Frank Neipel. gH was detected using custom-made polyclonal rabbit antibodies raised against peptides derived from KSHV gH (Genscript).

### Plasmids

The following plasmids were used in this study: pLenti CMV Blast DEST (706–1) (Ax203, a gift from Eric Campeau & Paul Kaufman, Addgene plasmid #17451); K8.1-OneStrep (OneStrep Tag: SAWSHPQFEKGGGSGGGSGGSAWSHPQFEK) in the Ax203 backbone (Ax336); pcDNA6aV5his encoding KSHV gH (Ax607); pcDNA3.1 encoding codon optimized KHSVgL (Ax252); pcDNA6aV5his encoding RRV gB (Ax223); VP16-Gal4 (Ax388)(49); The KSHV gH ectodomain (amino acids 22 through 704) N-terminally fused to the murine IgGkappa signal peptide and C-terminally fused to FcFcStrep (Strep tag:

SAWSHPQFEKGGGSGGGSGGGSWSHPQFEK) (Ax207)(19); the K8.1 ectodomain (amino acid 1 through 196) fused to FcFcStrep in the same backbone as Ax207 (Ax250); RRV 26-95 N-terminally fused to the murine IgGkappa signal peptide and C-terminally fused to FcFcStrep, same backbone as Ax207 (Ax176); RRV 26-95 R8.1 ectodomain (amino acid 26 through 237) N-terminally fused to the murine IgGkappa signal peptide and C-terminally fused to FcFcStrep in the same backbone as Ax207 (Ax243); pLenti CMV Blast DEST encoding a Gal4-driven TurboGFP-Luciferase fusion reporter gene (Ax526)(38).

### Infection assays and flow cytometry

For infection assay, cells were plated at 25 000 cells/well in 96-well plates. One day after plating, the cells were infected with the indicated amounts of virus. For HaCaT cells, spin infection was used. After adding the virus, HaCaT cells were centrifuged for 2 h at 800 g, 37°C. 48 h post infection, the cells were harvested by brief trypsinization, followed by adding 5% FCS in PBS to inhibit trypsin activity. Then, the cell suspension was transferred to an equal volume of PBS supplemented with 4% methanol-free formaldehyde (Carl Roth) for fixation. A minimum of 5 000 cells was analyzed per well for GFP expression on an ID7000 (Sony).

### Quantitative realtime-PCR-based viral genome copy number analysis and virus attachment assay

Concentrated virus samples were treated with DNAseI (0.1 units/μl) to remove non-encapsidated DNA (37°C, overnight). Subsequently, DNAseI was deactivated, and viral capsids were disrupted by heating the sample to 95°C for 30 minutes. Realtime-PCR (qPCR) was performed on a StepOne plus cycler (Thermo Fisher Scientific) in 20μl reactions using the Sensi FAST Probe Hi-ROX Kit (Bioline) (cycling conditions: 3min in initial denaturation at 95°C, 40 cycles 95°C for 10s and 60°C for 35s). All primer-probe sets were purchased from IDT as complete PrimeTime qPCR Assays (primer: probe ratio: 4:1). Samples were analyzed in technical duplicates. A series of six 10-fold dilutions of bacmid DNA was used as the standard for absolute quantification of viral genome copies based on qPCR of ORF59 of KSHV. HaCaT and SLK cells were seeded on the 12-well plate at 200 000 cells/well for virus attachment. After adding indicated amount of virus, cells are centrifuged for 30min at 4122g (4200rpm), at 4°C. After three washes with ice-cold PBS, genomic DNA was isolated using the ISOLATE II Genomic DNA Kit (Bioline) according to the manufacturer’s instructions. As described above, the ratio of viral DNA to cellular DNA as a measurement of the attached virus was determined by qPCR. Relative values of bound viral genomes to cellular DNA were calculated based on ΔCt values for viral genomic loci (ORF59 for KSHV) and a cellular genomic locus (CCR5).

### Protein production

Soluble K8.1-Fc, KSHV gH-Fc/gL, R8.1-Fc, and RRV gHΔ21-27-Fc protein were purified by Strep-Tactin chromatography from 293T cell culture supernatant. 293T cells were transfected by Polyethylenimine “Max” (PEI) (Polysciences) transfection with the respective expression plasmids. The protein-containing cell culture supernatant was passed over 0.5ml of a Strep-Tactin Superflow (IBA Lifescience) matrix in a gravity flow Omniprep column (BioRad). Bound protein was washed with approximately 50ml phosphate buffered saline pH 7.4 (PBS) and eluted in 1ml fractions with 3mM desthiobiotin (Sigma-Aldrich) in PBS. Protein-containing fractions were pooled, and buffer exchange to PBS via VivaSpin columns (Sartorius) was performed. Protein concentration was determined by absorbance at 280nm. Aliquots were frozen and stored at −80°C.

### Binding assay

Cells were detached by 0.25M EDTA in PBS and transferred to the falcon tube at 200 000 cells/tube. After discarding the supernatant, cells were fixed in 2% PFA for 10 min, followed by once washing with PBS. Subsequently, cells were blocked in 5% FCS in PBS for 1 h at 4°C and then incubated with 10 μg/m proteins in 5%FCS in PBS for 1 h at 4°C, followed by two washes with PBS and one wash with 5% FCS in PBS. Then, the cells were incubated with Alexa Fluor® 647 anti-human CD20 Antibody at a ratio of 1:200 for 45 min at 4°C in the dark, followed by three washes with PBS. Finally, the cells were stored in 1% PFA in PBS until flow cytometry analysis.

### Cell-cell fusion assay

SLK, HaCaT, A549, 293T target cells were stably transduced with a lentiviral construct encoding a Gal4 response element-driven TurboGFP-luciferase reporter. 293T effector cells were seeded in 96-well plates at 30 000 cells/well. One day after, 293T effector cells were transfected with a plasmid encoding the Gal4 DNA binding domain fused to the VP16 transactivation (VP16-Gal4), and the indicated viral glycoprotein combinations or a pcDNA empty vector control (VP16-Gal4: 31.25 ng/well, K8.1: 25ng/well, KSHV gH: 12.5 ng/well, KSHV gL: 62.5ng/well, RRV gB: 18.75 ng/well, pcDNA only: 118.75 ng/well) using PEI as described before. 24 h after transfection, the medium on 293T effector cells was removed entirely and exchanged to 100μl fresh DMEM supplemented with 10% FCS and 50μg/ml gentamicin. The target cells were re-suspended and counted, and 40 000 target cells were added to 293T effector cells in 100μl fresh DMEM supplemented with 10% FCS and 50 μg/ml gentamicin. Triplicate wells were analyzed for all target-effector combinations. After 48h, cells were washed once in PBS and lysed in 35 μl 1x Luciferase Cell culture lysis buffer (E1531, Promega) for 15min at room temperature and centrifuged for 10min at 4°C. 20 μl of each cell lysate was used to measure luciferase activity using the Beetle-Juice Luciferase Assay according to the manufacturer’s instructions on a Biotek Synergy 2 plate reader.

## ACKNOWLEDGEMENTS

This work was supported by grants to ASH from the Deutsche Forschungsgemeinschaft (www.dfg.de, HA 6013/4-1 and HA 6013/10-1) and from the Wilhelm-Sander-Foundation (www.wilhelmsander-stiftung.de, project 2019.027.1). We would like to thank Frank Neipel, Scott Wong, Klaus Korn, and Stefan Pöhlmann for sharing reagents and resources.

## REFERENCES

1. Cesarman E, Chang Y, Moore PS, Said JW, Knowles DM. 1995. Kaposi’s sarcoma-associated herpesvirus-like DNA sequences in AIDS-related body-cavity-based lymphomas. N Engl J Med 1:1186–p.

2. Chang Y, Cesarman E, Pessin MS, Lee F, Culpepper J, Knowles DM, Moore PS. 1994. Identification of herpesvirus-like DNA sequences in AIDS-associated Kaposi’s sarcoma. Science 1:1865–1869.

3. Parravicini C, Corbellino M, Paulli M, Magrini U, Lazzarino M, Moore PS, Chang Y. 1997. Expression of a virus-derived cytokine, KSHV vIL-6, in HIV-seronegative Castleman’s disease. Am J Pathol 1:1517–1522.

4. Soulier J, Grollet L, Oksenhendler E, Cacoub P, Cazals-Hatem D, Babinet P, d’Agay MF, Clauvel JP, Raphael M, Degos L. 1995. Kaposi’s sarcoma-associated herpesvirus-like DNA sequences in multicentric Castleman’s disease. Blood 1:1276–1280.

5. Wen KW, Wang L, Menke JR, Damania B. 2022. Cancers associated with human gammaherpesviruses. The FEBS Journal 1:7631–7669.

6. Chen Q, Chen J, Li Y, Liu D, Zeng Y, Tian Z, Yunus A, Yang Y, Lu J, Song X, Yuan Y. 2021. Kaposi’s sarcoma herpesvirus is associated with osteosarcoma in Xinjiang populations. PNAS 118.

7. Goncalves PH, Ziegelbauer J, Uldrick TS, Yarchoan R. 2017. Kaposi sarcoma herpesvirus-associated cancers and related diseases. Current Opinion in HIV and AIDS 1:47–56.

8. Cesarman E, Damania B, Krown SE, Martin J, Bower M, Whitby D. 2019. Kaposi sarcoma. Nat Rev Dis Primers 1:1–21.

9. Yarchoan R, Uldrick TS. 2018. HIV-Associated Cancers and Related Diseases. N Engl J Med 1:1029–1041.

10. Bray F, Ferlay J, Soerjomataram I, Siegel RL, Torre LA, Jemal A. 2018. Global cancer statistics 1: GLOBOCAN estimates of incidence and mortality worldwide for 36 cancers in 185 countries. CA Cancer J Clin https://doi.org/10.3322/caac.21492.

11. Connolly SA, Jardetzky TS, Longnecker R. 2021. The structural basis of herpesvirus entry. Nat Rev Microbiol 1:110–121.

12. Garrigues HJ, DeMaster LK, Rubinchikova YE, Rose TM. 2014. KSHV attachment and entry are dependent on αVβ3 integrin localized to specific cell surface microdomains and do not correlate with the presence of heparan sulfate. Virology 1:118–133.

13. Garrigues HJ, Rubinchikova YE, DiPersio CM, Rose TM. 2008. Integrin {alpha}V 3 Binds to the RGD Motif of Glycoprotein B of Kaposi’s Sarcoma-Associated Herpesvirus and Functions as an RGD-Dependent Entry Receptor. J Virol 1:1570–1580.

14. Akula SM, Pramod NP, Wang F-Z, Chandran B. 2002. Integrin α3β1 (CD 49c/29) Is a Cellular Receptor for Kaposi’s Sarcoma-Associated Herpesvirus (KSHV/HHV-8) Entry into the Target Cells. Cell 1:407–419.

15. Veettil MV, Sadagopan S, Sharma-Walia N, Wang F-Z, Raghu H, Varga L, Chandran B. 2008. Kaposi’s Sarcoma-Associated Herpesvirus Forms a Multimolecular Complex of Integrins (αVβ5, αVβ3, and α3β1) and CD98-xCT during Infection of Human Dermal Microvascular Endothelial Cells, and CD98-xCT Is Essential for the Postentry Stage of Infection. Journal of Virology 1:12126–12144.

16. Chen J, Sathiyamoorthy K, Zhang X, Schaller S, Perez White BE, Jardetzky TS, Longnecker R. 2018. Ephrin receptor A2 is a functional entry receptor for Epstein-Barr virus. Nat Microbiol 1:172–180.

17. Chen J, Zhang X, Schaller S, Jardetzky TS, Longnecker R. 2019. Ephrin Receptor A4 is a New Kaposi’s Sarcoma-Associated Herpesvirus Virus Entry Receptor. mBio 1:e02892–18.

18. Großkopf AK, Schlagowski S, Fricke T, Ensser A, Desrosiers RC, Hahn AS. 2021. Plxdc family members are novel receptors for the rhesus monkey rhadinovirus (RRV). PLoS Pathog 1:e1008979.

19. Hahn AS, Kaufmann JK, Wies E, Naschberger E, Panteleev-Ivlev J, Schmidt K, Holzer A, Schmidt M, Chen J, König S, Ensser A, Myoung J, Brockmeyer NH, Stürzl M, Fleckenstein B, Neipel F. 2012. The ephrin receptor tyrosine kinase A2 is a cellular receptor for Kaposi’s sarcoma–associated herpesvirus. Nat Med 1:961–966.

20. Chakraborty S, Valiyaveettil M, Sadagopan S, Paudel N, Chandran B. 2011. c-Cbl-Mediated Selective Virus-Receptor Translocations into Lipid Rafts Regulate Productive Kaposi’s Sarcoma-Associated Herpesvirus Infection in Endothelial Cells. J Virol 1:12410–12430.

21. Dutta D, Chakraborty S, Bandyopadhyay C, Valiya Veettil M, Ansari MA, Singh VV, Chandran B. 2013. EphrinA2 Regulates Clathrin Mediated KSHV Endocytosis in Fibroblast Cells by Coordinating Integrin-Associated Signaling and c-Cbl Directed Polyubiquitination. PLoS Pathog 1:e1003510.

22. Kaleeba JAR, Berger EA. 2006. Kaposi’s Sarcoma-Associated Herpesvirus Fusion-Entry Receptor: Cystine Transporter xCT. Science 1:1921–1924.

23. Raab MS, Albrecht JC, Birkmann A, Yağuboğlu S, Lang D, Fleckenstein B, Neipel F. 1998. The immunogenic glycoprotein gp35-37 of human herpesvirus 8 is encoded by open reading frame K8.1. J Virol 1:6725–6731.

24. Birkmann A, Mahr K, Ensser A, Yağuboğlu S, Titgemeyer F, Fleckenstein B, Neipel F. 2001. Cell Surface Heparan Sulfate Is a Receptor for Human Herpesvirusl18 and Interacts with Envelope Glycoprotein K8.1. Journal of Virology 1:11583–11593.

25. Dollery SJ, Santiago-Crespo RJ, Chatterjee D, Berger EA. 2018. Glycoprotein K8.1A of Kaposi’s sarcoma-associated herpesvirus is a critical B cell tropism determinant, independent of its heparan sulfate binding activity. J Virol https://doi.org/10.1128/JVI.01876-18.

26. Haan KM, Kyeong Lee S, Longnecker R. 2001. Different Functional Domains in the Cytoplasmic Tail of Glycoprotein B Are Involved in Epstein–Barr Virus-Induced Membrane Fusion. Virology 1:106–114.

27. Arias C, Weisburd B, Stern-Ginossar N, Mercier A, Madrid AS, Bellare P, Holdorf M, Weissman JS, Ganem D. 2014. KSHV 2.1: a comprehensive annotation of the Kaposi’s sarcoma-associated herpesvirus genome using next-generation sequencing reveals novel genomic and functional features. PLoS Pathog 1:e1003847.

28. Stürzl M, Gaus D, Dirks WG, Ganem D, Jochmann R. 2013. Kaposi’s sarcoma-derived cell line SLK is not of endothelial origin, but is a contaminant from a known renal carcinoma cell line. Int J Cancer 1:1954–1958.

29. Deyrieux AF, Wilson VG. 2007. In vitro culture conditions to study keratinocyte differentiation using the HaCaT cell line. Cytotechnology 1:77–83.

30. Reed JA, Nador RG, Spaulding D, Tani Y, Cesarman E, Knowles DM. 1998. Demonstration of Kaposi’s sarcoma-associated herpes virus cyclin D homolog in cutaneous Kaposi’s sarcoma by colorimetric in situ hybridization using a catalyzed signal amplification system. Blood 1:3825–3832.

31. Cerimele F, Curreli F, Ely S, Friedman-Kien AE, Cesarman E, Flore O. 2001. Kaposi’s sarcoma-associated herpesvirus can productively infect primary human keratinocytes and alter their growth properties. J Virol 1:2435–2443.

32. Foreman KE, Bacon PE, Hsi ED, Nickoloff BJ. 1997. In situ polymerase chain reaction-based localization studies support role of human herpesvirus-8 as the cause of two AIDS-related neoplasms: Kaposi’s sarcoma and body cavity lymphoma. J Clin Invest 1:2971–2978.

33. Boukamp P, Petrussevska RT, Breitkreutz D, Hornung J, Markham A, Fusenig NE. 1988. Normal keratinization in a spontaneously immortalized aneuploid human keratinocyte cell line. J Cell Biol 1:761–771.

34. Großkopf AK, Schlagowski S, Hörnich BF, Fricke T, Desrosiers RC, Hahn AS. 2019. EphA7 Functions as Receptor on BJAB Cells for Cell-to-Cell Transmission of the Kaposi’s Sarcoma-Associated Herpesvirus and for Cell-Free Infection by the Related Rhesus Monkey Rhadinovirus. J Virol 1:e00064–19.

35. Dollery SJ, Santiago-Crespo RJ, Kardava L, Moir S, Berger EA. 2014. Efficient infection of a human B cell line with cell-free Kaposi’s sarcoma-associated herpesvirus. J Virol 1:1748–1757.

36. Myoung J, Ganem D. 2011. Infection of Lymphoblastoid Cell Lines by Kaposi’s Sarcoma-Associated Herpesvirus: Critical Role of Cell-Associated Virus. Journal of Virology 1:9767–9777.

37. Hahn A, Birkmann A, Wies E, Dorer D, Mahr K, Stürzl M, Titgemeyer F, Neipel F. 2009. Kaposi’s sarcoma-associated herpesvirus gH/gL: glycoprotein export and interaction with cellular receptors. J Virol 1:396–407.

38. Hörnich BF, Großkopf AK, Dcosta CJ, Schlagowski S, Hahn AS. 2021. Interferon-Induced Transmembrane Proteins Inhibit Infection by the Kaposi’s Sarcoma-Associated Herpesvirus and the Related Rhesus Monkey Rhadinovirus in a Cell-Specific Manner. mBio 1:e0211321.

39. Fricke T, Großkopf AK, Ensser A, Backovic M, Hahn AS. 2022. Antibodies Targeting KSHV gH/gL Reveal Distinct Neutralization Mechanisms. 3. Viruses 1:541.

40. Colombo I, Sangiovanni E, Maggio R, Mattozzi C, Zava S, Corbett Y, Fumagalli M, Carlino C, Corsetto PA, Scaccabarozzi D, Calvieri S, Gismondi A, Taramelli D, Dell’Agli M. 2017. HaCaT Cells as a Reliable In Vitro Differentiation Model to Dissect the Inflammatory/Repair Response of Human Keratinocytes. Mediators Inflamm 1:7435621.

41. Hayes AJ, Melrose J. 2023. HS, an Ancient Molecular Recognition and Information Storage Glycosaminoglycan, Equips HS-Proteoglycans with Diverse Matrix and Cell-Interactive Properties Operative in Tissue Development and Tissue Function in Health and Disease. 2. International Journal of Molecular Sciences 1:1148.

42. Tanner J, Weis J, Fearon D, Whang Y, Kieff E. 1987. Epstein-Barr virus gp350/220 binding to the B lymphocyte C3d receptor mediates adsorption, capping, and endocytosis. Cell 1:203–213.

43. Hahn AS, Desrosiers RC. 2013. Rhesus Monkey Rhadinovirus Uses Eph Family Receptors for Entry into B Cells and Endothelial Cells but Not Fibroblasts. PLOS Pathogens 1:e1003360.

44. Brulois KF, Chang H, Lee AS-Y, Ensser A, Wong L-Y, Toth Z, Lee SH, Lee H-R, Myoung J, Ganem D, Oh T-K, Kim JF, Gao S-J, Jung JU. 2012. Construction and Manipulation of a New Kaposi’s Sarcoma-Associated Herpesvirus Bacterial Artificial Chromosome Clone. Journal of Virology 1:9708–9720.

45. Myoung J, Ganem D. 2011. Generation of a doxycycline-inducible KSHV producer cell line of endothelial origin: Maintenance of tight latency with efficient reactivation upon induction. Journal of Virological Methods 1:12–21.

46. Tischer BK, von Einem J, Kaufer B, Osterrieder N. 2006. Two-step red-mediated recombination for versatile high-efficiency markerless DNA manipulation in Escherichia coli. BioTechniques 1:191–197.

47. Großkopf AK, Ensser A, Neipel F, Jungnickl D, Schlagowski S, Desrosiers RC, Hahn AS. 2018. A conserved Eph family receptor-binding motif on the gH/gL complex of Kaposi’s sarcoma-associated herpesvirus and rhesus monkey rhadinovirus. PLoS Pathog 1:e1006912.

48. Lang D, Birkmann A, Neipel F, Hinderer W, Rothe M, Ernst M, Sonneborn HH. 2000. Generation of monoclonal antibodies directed against the immunogenic glycoprotein K8.1 of human herpesvirus 8. Hybridoma 1:287–295.

49. Hörnich BF, Großkopf AK, Schlagowski S, Tenbusch M, Kleine-Weber H, Neipel F, Stahl-Hennig C, Hahn AS. 2021. SARS-CoV-2 and SARS-CoV Spike-Mediated Cell-Cell Fusion Differ in Their Requirements for Receptor Expression and Proteolytic Activation. J Virol 1:e00002–21.

